# Indomethacin has a potent antiviral activity against SARS CoV-2 in vitro and canine coronavirus in vivo

**DOI:** 10.1101/2020.04.01.017624

**Authors:** Tianhong Xu, Xuejuan Gao, Zengbin Wu, Douglas W. Selinger, Zichen Zhou

## Abstract

**Background:** The outbreak of SARS CoV-2 has caused ever-increasing attention and public panic all over the world. Currently, there is no specific treatment against the SARS CoV-2. Therefore, identifying effective antiviral agents to combat the disease is urgently needed. Previous studies found that indomethacin has the ability to inhibit the replication of several unrelated DNA and RNA viruses, including SARS-CoV.

**Methods:** SARS CoV-2 pseudovirus-infected African green monkey kidney VERO E6 cells treated with different concentrations of indomethacin or aspirin at 48 hours post infection (p.i). The level of cell infection was determined by luciferase activity. Anti-coronavirus efficacy in vivo was confirmed by evaluating the time of recovery in canine coronavirus (CCV) infected dogs treated orally with 1mg/kg body weight indomethacin.

**Results:** We found that indomethacin has a directly and potently antiviral activity against the SARS CoV-2 pseudovirus (reduce relative light unit to zero). In CCV-infected dogs, recovery occurred significantly sooner with symptomatic treatment + oral indomethacin (1 mg/kg body weight) daily treatments than with symptomatic treatment + ribavirin (10-15 mg/kg body weight) daily treatments (P =0.0031), but was not significantly different from that with symptomatic treatment + anti-canine coronavirus serum + canine hemoglobin + canine blood immunoglobulin + interferon treatments (P =0.7784).

**Conclusion:** The results identify indomethacin as a potent inhibitor of SARS CoV-2.

## Introduction

Since 12 December 2019, there has been an outbreak of Acute Respiratory Syndrome caused by a new coronavirus (SARS CoV-2). The disease caused by the virus was named Coronavirus Disease 19 (COVID-19). As of March 20, 2020, SARS CoV-2 has been reported in 76 countries and 308,613 cases and 13,844 deaths have been confirmed, with an estimated mortality risk of 4%. Common symptoms at onset of illness were fever, cough, myalgia and fatigue; fewer common symptoms were sputum production, headache, haemoptysis, and diarrhea. Complications included acute respiratory distress syndrome, acute cardiac injury ^1^.

It is currently believed that SARS CoV-2 may be transmitted from bats to humans^2^. Angiotensin-converting enzyme 2 (ACE2) is the main host cell receptor of SARS CoV-2 and plays a crucial role in the entry of virus into the cell to cause infection^3-5^. High ACE2 expression was identified in type II alveolar cells (AT2) of lung, esophagus upper and stratified epithelial cells, absorptive enterocytes from ileum and colon, and bladder urothelial cells^6-8^.

New interventions are likely to require years to develop. Given the urgency of the SARS CoV-2 outbreak, we focus here on the potential to repurpose existing drugs approved for treating infections caused by RNA virus, based on therapeutic experience with two other infections caused by human coronaviruses: severe acute respiratory syndrome (SARS) and Middle East respiratory syndrome (MERS)^9^. Cyclooxygenases (COXs) play a significant role in many different viral infections with respect to replication and pathogenesis^10^. Cyclopentone COX inhibitor indomethacin has been widely used in the clinic for its potent anti-inflammatory and analgesic properties^11^. In addition to the anti-inflammatory/analgesic action, indomethacin has been known to also possess antiviral properties, including canine coronavirus (CCV) and SARS-CoV^12-14^.

CCV has certain sequence homology with SARS CoV-2^12^. CCV-infection in dogs are common in winter and spread rapidly. Puppies have higher morbidity and mortality than adult dogs^15^. The main symptoms of dogs with CCV-infection are those of the digestive system, such as vomiting and diarrhea, and sometimes respiratory symptoms^16^.

In the present report we described how indomethacin was found to potently inhibit SARS CoV-2 *in vitro*, and efficiently treat CCV-infected dogs.

## Methods

### Cell culture and treatment

VERO E6 cells were grown at 37°C in a 5% CO_2_ humidified atmosphere in Modified Eagle Medium (MEM), supplemented with 10% fetal calf serum (FCS). Indomethacin was dissolved in absolute ethanol at a concentration of (0, 0.1, 1, 5, 10, 50, 100, 500 µM) and diluted in culture medium immediately before use. Unless otherwise specified, indomethacin and Aspirin were added immediately after the 1 h adsorption period and maintained in the medium for the duration of the experiment. Controls received equal amounts of appropriate diluent Cell viability was determined by 3-(4,5-dimethylthiazol-2-yl)-2,5-diphenyl-tetrazolium bromide (MTT) to MTT formazan conversion assay.

### Coronavirus infection and titration

The cDNA of the SARS CoV-2 (SARS CoV-2 S,GenBank: MN908947.3). The SARS CoV-2 pseudotyped virus HIV/SARS was produced by a similar method which was described previously^17^. Briefly, 10 µg pNL4-3-Luc-R-E-pro^18^ and 10 µg pTSh were co-transfected into VERO E6 cells in 10 cm dishes. Supernatants were harvested 48 h later and used in infection assays. The pseudotyped virus was purified by ultracentrifugation through a 20% sucrose cushion at 50,000 g for 90 min, resuspended in 100 µl PBS. The supernatant containing 5 ng pseudotyped virus (p24) was used to infect cells in 24-well plates (4–8×10^4^ cells/well), and treated with different concentrations (0, 0.1, 1, 5, 10, 50, 100, 500 µM) of indomethacin or Aspirin after the 1 h adsorption period. The cells were lysed at 48 h post infection (p.i). Twenty microliters of lysate was tested for luciferase activity by the addition of 50 µl of luciferase substrate and measured for 10 s in a Wallac Multilabel 1450 Counter. After incubation at 37°C for 30 min, 100 µl mixtures were added to VERO E6 cells in 96-well plates. The cells were lysed at 48 h p.i and tested for luciferase activity as described above^19^.

### Animal maintenance

The enrolled dogs were confirmed for diagnosis of CCV infection with a Canine Coronavirus test kit and between 2-3 days of the onset of symptoms. We enrolled 26 dogs which were 3-6 months old and medium-sized. There were several types of these dogs, including poodles and Pekingese dogs. Dogs were grouped into three populations based on the treatment scheme as follows: (1) symptomatic treatment + ribavirin (10-15 mg/kg body weight) daily, 8 dogs; (2) symptomatic treatment + anti-canine coronavirus serum + canine hemoglobin + canine blood immunoglobulin + interferon, 9 dogs; (3) symptomatic treatment + oral indomethacin (1 mg/kg body weight) daily^20^, 9 dogs. The time of recovery was determined by the disappearance of symptoms and a negative diagnosis by the canine coronavirus test kit. Humane standards were adhered to for all assessments. The study procedure was approved by Animal Ethics Committee of United Animal Hospital.

### Statistical analysis

In virus titration experiments, statistical analysis was performed using Student’s t-test for unpaired data. Data were expressed as the mean ± SD and P-values of <0.05 were considered significant.

## Results

### 1. Indomethacin is a potent inhibitor of SARS CoV-2 *in vitro* replication

To investigate the effect of indomethacin on SARS CoV-2 replication, African green monkey kidney VERO E6 cells were infected with SARS CoV-2 and treated with different concentrations (0, 0.1, 1, 5, 10, 50, 100, 500 µM) of indomethacin or Aspirin after the 1 h adsorption period. The level of cell infection was determined by luciferase activity at 48 h p.i. Surprisingly, indomethacin was found to possess a remarkable antiviral activity, reducing viral particle production dose-dependently with an IC50 (inhibitory concentration 50%) of 1 µM, and selective index of 500, and caused a dramatic reduction relative light unit to zero at 48 h p.i, in VERO E6 cells (Figure 1A). Oral administration of 50 mg of indomethacin generates a peak plasma concentration of the drug ranging from 7µM to 11µM, a concentration which is 10 times of the drug’s IC50^21^. These concentrations of indomethacin did not affect VERO E6 cell proliferation (Figure 1B) indicating that even at 500 µM indomethacin still does not show any cytotoxicity to VERO E6 cell.

**Figure 1.**
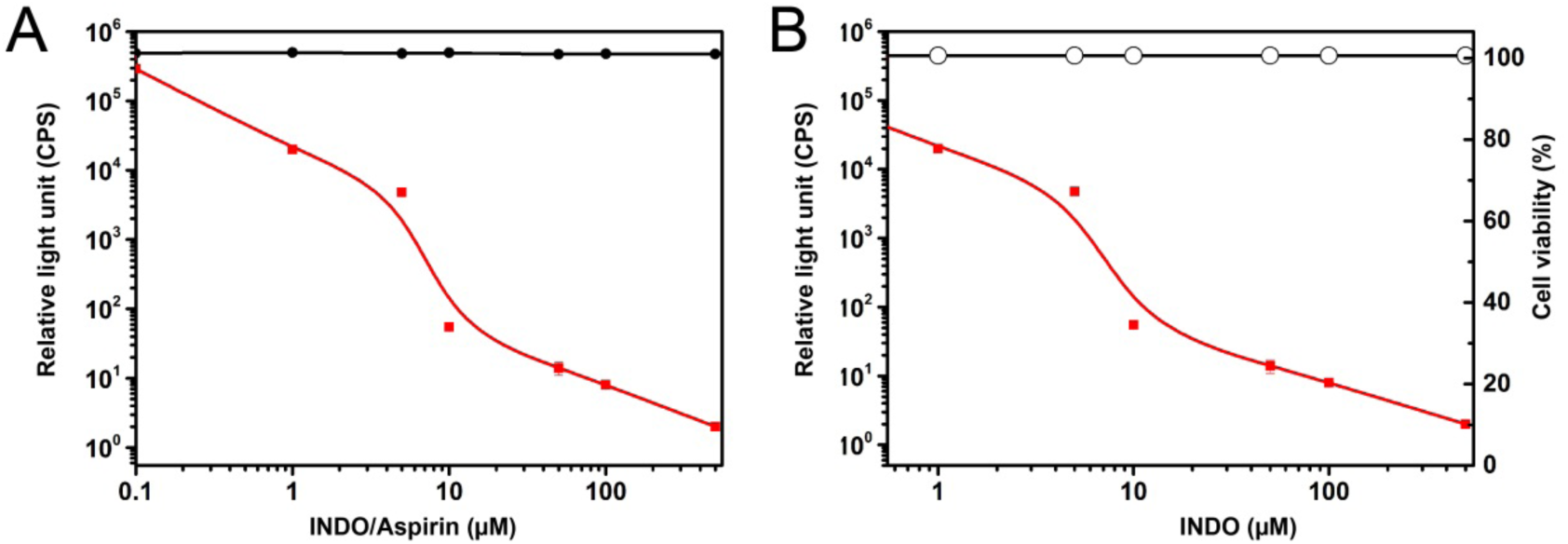
Indomethacin is a potent inhibitor of SARS CoV-2 in vitro replication. (A). VERO E6 cells were infected with SARS CoV-2 and treated with different concentrations of the non-steroidal anti-inflammatory drugs (NSAIDs) indomethacin (INDO, red line) (0, 0.1, 1, 5, 10, 50, 100, 500 µM) or Aspirin (black line) (0, 0.1, 1, 5, 10, 50, 100, 500 µM) after the 1 h adsorption period. The level of cell infection was determined by luciferase activity in the supernatant of infected cells at 48 h p.i. (B). Cell viability (black line) determined by MTT assay in uninfected indomethacin-treated cells is expressed as percentage of MTT conversion in untreated control, and (red line) indicates relative light unit treated with as (A). Data expressed in relative light unit represent the mean ± SD of triplicate samples from a representative experiment of three with similar results. P <0.05 indicates a significant difference.

### 2. Antiviral activity of indomethacin in CCV-infected dogs

In order to evaluate whether indomethacin could also be effective against CCV *in vivo*, we analyzed the rate of recovery in dogs with treatment schemes as follows: (1) symptomatic treatment + ribavirin (10-15 mg/kg body weight) daily; (2) symptomatic treatment + anti-canine coronavirus serum + canine hemoglobin + canine blood immunoglobulin + interferon; (3) symptomatic treatment + oral indomethacin (1 mg/kg body weight) daily. The results showed that recovery occurred significantly sooner with the (3) treatments than with the (1) treatments (P =0.0031, Fig.2A), but was not significantly different from that with the (2) treatments (P =0.7784, Fig.2B). Moreover, in the group of with the (1) treatment scheme were 8 dogs, and 3 died (3/8); with the (2) scheme were 9 dogs, and 1 died (1/9); with the (3) scheme were all survived from this disease (9 dogs).

**Figure 2.**
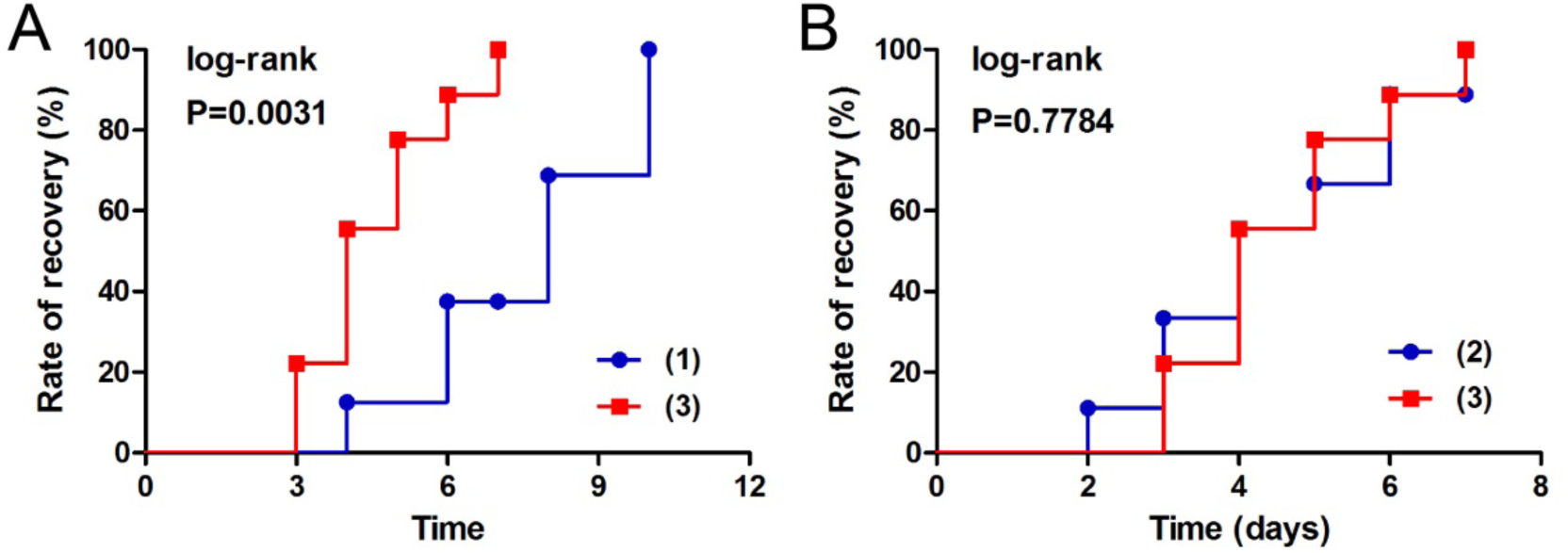
Antiviral activity of indomethacin in CCV-infected dogs. This figure shows the rate of recovery in CCV-infected dogs with the treatment schemes as follows: (1) symptomatic treatment + ribavirin (10-15 mg/kg body weight) daily; (2) symptomatic treatment + anti-canine coronavirus serum + canine hemoglobin + canine blood immunoglobulin + interferon; (3) symptomatic treatment + oral indomethacin (1 mg/kg body weight) daily. The time of recovery was determined by the symptoms disappeared and the diagnosis of CCV test paper turned negative. Both in (A) and (B) red line indicates treatment with the (3) scheme. Blue line indicates treatment with the (1) scheme in (A), however indicate treatment with the (2) scheme in (B). P <0.05 indicates a significant difference.

## Discussion

Given the urgency of the SARS CoV-2 outbreak, we focus here on the potential to repurpose existing agents approved or in development for treating infections caused by HIV, hepatitis B virus (HBV), hepatitis C virus (HCV) and influenza^9,22^. Indomethacin has been known to possess antiviral properties^12-14^, and used for a long time as a potent anti-inflammatory drug, acting by blocking the activity of cyclooxygenase-1 and −2 (COXs), activity and inhibiting pro-inflammatory prostaglandin synthesis^23^. Indomethacin reduces the synthesis of prostaglandin (PG) by inhibiting cyclooxygenase, and inhibiting the formation of painful nerve impulses in inflammatory tissues thus suppressing the inflammatory response^10^.

In our experiments, we tested the COX inhibitors indomethacin and aspirin. Our results show that indomethacin has potent, direct antiviral activity against the SARS CoV-2 (reduce the relative light unit to zero) in VERO E6 cells (Figure 1A), and does not show cytotoxicity to VERO E6 cells (Figure 1B). The current study, and reported by others, suggest that indomethacin has strong anti-Coronavirus activity *in vitro*^12,13^, including SARS COV-2, while aspirin does not. This suggests that indomethacin’s antiviral activity is not achieved by some mechanisms other than COX inhibition.

In our study of CCV-infected dogs, the results show that in items of anti-CCV efficacy, indomethacin can achieve a similar efficacy as treatment with anti-canine coronavirus serum, canine hemoglobin, canine blood immunoglobulin and interferon treatments effect (P =0.7784, Fig.2B), and superior efficacy than the treatment with ribavirin (P =0.0031, Fig.2A). These findings were consistent with other report and revealed that indomethacin also shows significant anti-CCV *in vivo*^12^. Its mechanism, however, remains largely unknown.

Although the underlying mechanism requires further investigation, the results of indomethacin anti-SARS-CoV show that indomethacin does not affect virus infectivity, binding or entry into target cells, but acts early on the coronavirus replication cycle, selectively blocking viral RNA synthesis^12^. The only clear indomethacin antiviral mechanism is that, instead, triggers a cellular antiviral defense mechanism by rapidly and effectively activating PKR in an interferon- and dsRNA-independent manner^13^. PKR plays a critical role in the antiviral defense mechanism of the host, acting as a sensor of virus replication and^24^, upon activation, leading to eIF2α phosphorylation and block of protein synthesis in virally infected cells^25^.

To investigate possible antiviral mechanisms not related to COX inhibition, we utilized Plex Research’s AI search engine platform^26^. A Plex analysis of indomethacin identified additional targets that are directly modulated by indomethacin or its close analogs: aldo-keto reductase (AKR1C3, AKR1C4, AKR1C2)^27-29^, aldose reductase (AKR1B1)^30^, peroxisome proliferator activated receptor gamma (PPARG)^31-33^, and cannabinoid receptor 2 (CB2)^34^. Of these, only AKR1B1 and PPARG are expressed in VERO cells^35^.

There are few reports about the role of aldo-keto reductases in inflammation. Aldo-keto reductase was found to reduce peroxidation derived lipid aldehydes, such as 4-hydroxytrans-2-nonenal and their glutathione conjugates, to corresponding alcohols, 1,4-dihydroxy-nonene and glutathionyl-1,4-dihydroxynonene, respectively, which take part in the inflammatory signaling^36^. Inhibition of aldose reductase was found to significantly prevent the transfer of the inflammatory signals induced by multiple factors including allergens, cytokines, growth factors, endotoxins, high glucose, and autoimmune reactions at the cellular level as well as in animal models^37^.

It has been demonstrated that PPARG agonists inhibit the production of pro-inflammatory cytokines, such as TNF-α, IL-1β, and IL-6^38^. Macrophages loss of PPARG leads to altered differentiation kinetics^39^. PPARG has been described to be important for resolving inflammation and maintaining homeostasis.

Cannabinoids receptor 2 (CB2), which is influences the immune system, viral pathogenesis, and viral replication. The anti-inflammatory and immunoregulatory action of cannabinoids are CB2 dependent^40^. Activated CB2 receptors play a significant role in leukocyte and endothelial migration, activation, and interaction, which is related to the anti-inflammatory and immunomodulatory effects of CB2^41^. CB2 receptors suppress chemokine induced chemotaxis of neutrophils, lymphocytes, macrophages, monocytes, and microglia by inhibiting leukocyte migration mediated by RhoA activation^42,43^. In addition, CB2 can also inhibit the recruitment of neutrophils^44^. Therefore, CB2 plays a crucial role in maintaining immune homeostasis and controlling the magnitude of the immune response through negative regulation.

IL-6 is a soluble mediator with a pleiotropic effect on inflammation and immune response^45^. It is generally known as endogenous pyrogens which can induce fever. It was shown that an increase in intrahypothalamic concentrations of IL-6 and TNF-like activity during LPS-induces fever^46^. A recent report indicates that indomethacin can reduce the cytokine-mediated IL-6 release by reducing LPS^47^. COVID-19 patients show a “cytokine storm” phenomenon, while IL-6 in the body is significantly increased^47,48^. The latest treatment guidelines for COVID-19 include the reduction of concentration of IL-6.

During the preparation of this manuscript, a preliminary report emerged describing a proteomic analysis of interactions between SARS CoV-2 viral proteins and 332 human proteins^49^. In this report, the viral NSP7 protein was found to interact with human PTGES2. PTGES2 is expressed in VERO cells^50^ and is inhibited by indomethacin (IC_50_ = 750 nM)^51^. It is also inhibited by aspirin, however much less potently (IC_50_ = 35 μM)^51^. NSP7, together with NSP8, forms the viral primase complex, which is part of the viral RNA polymerase machinery^52,53^. Inhibition of NSP7/NSP8 complex function through inhibition of PTGES2 would be consistent with the selective blockage of viral RNA synthesis observed after indomethacin treatment of SARS-CoV^12^.

Although the mechanisms underlying the antiviral activity of indomethacin against coronavirus infection require further investigation, our results herein show that indomethacin inhibits SARS CoV-2 both *in vitro* and *in vivo*.

Previously, we have used nebulized indomethacin to treat patients with bronchorrhea or excessive phlegm caused by bronchioloalveolar cell carcinoma or COPD and found that indomethacin can dramatically reduce the bronchorrhea sputum and abate dyspnea. Due to patients with COVID-19 often exhibit excessive phlegm which may causes severe dyspnea, indomethacin not only directly inhibit virus as our current study identified, but it also may reduce sputum secretion and relieve symptoms.

NSAIDs are commonly used as antipyretics and/or analgesics in treating Community-acquired pneumonia. In some survey or observational study, up to half of the patients diagnosed with CAP were treated with NSAIDs^54-56^. Among these NSAIDs, indomethacin is used less frequently than other paracetamol, fenbufen, ibuprofen or aspirin, mainly due to the relatively frequent gastrointestinal side effects of oral indomethacin, but indomethacin rarely causes serious side effects. Given the fact that indomethacin has been widely used in the clinic for several decades and its safety profile is well defined^57^, we suggest that indomethacin should be assessed further in a prospective clinical study as a treatment of COVID-19 patients.

## Acknowledgements

We thank Professor Tao ShenCe from Shanghai Center for System Biomedicine for providing SARS COV-2 coding region plasmid. We thank Prof. M. Gabriella Santoro from University of Rome Tor Vergata for valuable discussion, and we thank Vic Lee and Virtus Inspire Ventures providing venture fund supporting this work.

## References

1. Huang C, Wang Y, Li X, et al. Clinical features of patients infected with 2019 novel coronavirus in Wuhan, China. Lancet. 2020;395(10223):497–506.

2. Zhou P, Yang X-L, Wang X-G, et al. A pneumonia outbreak associated with a new coronavirus of probable bat origin. Nature. 2020:10.1038/s41586-41020-42012-41587.

3. Letko M, Marzi A, Munster V. Functional assessment of cell entry and receptor usage for SARS-CoV-2 and other lineage B betacoronaviruses. Nat Microbiol. 2020.

4. Xu H, Zhong L, Deng J, et al. High expression of ACE2 receptor of 2019-nCoV on the epithelial cells of oral mucosa. Int J Oral Sci. 2020;12(1):8.

5. Cao Y, Li L, Feng Z, et al. Comparative genetic analysis of the novel coronavirus (2019-nCoV/SARS-CoV-2) receptor ACE2 in different populations. Cell Discovery. 2020;6(1).

6. Xin Zou Kcjzphjhzh. The single-cell RNA-seq data analysis on the receptor ACE2 expression reveals the potential risk of different human organs vulnerable to Wuhan 2019-nCoV infection. Frontiers of Medicine. 0.

7. Zhang H, Kang Z, Gong H, et al. The digestive system is a potential route of 2019-nCov infection: a bioinformatics analysis based on single-cell transcriptomes. bioRxiv. 2020: 2020.2001.2030.927806.

8. Zhao Y, Zhao Z, Wang Y, Zhou Y, Ma Y, Zuo W. Single-cell RNA expression profiling of ACE2, the putative receptor of Wuhan 2019-nCov. bioRxiv. 2020:2020.2001.2026.919985.

9. Li G, De Clercq E. Therapeutic options for the 2019 novel coronavirus (2019-nCoV). Nat Rev Drug Discov. 2020;19(3):149–150.

10. Raaben M, Einerhand AWC, Taminiau LJA, et al. Cyclooxygenase activity is important for efficient replication of mouse hepatitis virus at an early stage of infection. Virology Journal. 2007;4(1).

11. Vane JR, Botting RM. Mechanism of action of nonsteroidal anti-inflammatory drugs. Am J Med. 1998;104(3A):2S–22S.

12. Amici C, Di Caro A, Ciucci A, et al. Indomethacin has a potent antiviral activity against SARS coronavirus. Antivir Ther. 2006;11(8):1021–1030.

13. Amici C, La Frazia S, Brunelli C, Balsamo M, Angelini M, Santoro MG. Inhibition of viral protein translation by indomethacin in vesicular stomatitis virus infection: role of eIF2alpha kinase PKR. Cell Microbiol. 2015;17(9):1391–1404.

14. Rossen JWA, Bouma J, Raatgeep RHC, Büller HA, Einerhand AWC. Inhibition of cyclooxygenase activity reduces rotavirus infection at a postbinding step. J Virol. 2004;78(18):9721–9730.

15. Buonavoglia C, Decaro N, Martella V, et al. Canine coronavirus highly pathogenic for dogs. Emerg Infect Dis. 2006;12(3):492–494.

16. van Nguyen D, Terada Y, Minami S, et al. Characterization of canine coronavirus spread among domestic dogs in Vietnam. J Vet Med Sci. 2017;79(2):343–349.

17. Deng H, Liu R Fau - Ellmeier W, Ellmeier W Fau - Choe S, et al. Identification of a major co-receptor for primary isolates of HIV-1. (0028-0836 (Print)).

18. Connor RI, Chen Bk Fau - Choe S, Choe S Fau - Landau NR, Landau NR. Vpr is required for efficient replication of human immunodeficiency virus type-1 in mononuclear phagocytes. (0042-6822 (Print)).

19. Nie Y, Wang P Fau - Shi X, Shi X Fau - Wang G, et al. Highly infectious SARS-CoV pseudotyped virus reveals the cell tropism and its correlation with receptor expression. (0006-291X (Print)).

20. Arai I, Mao Gp Fau - Otani K, Otani K Fau - Konno S, Konno S Fau - Kikuchi S, Kikuchi S Fau - Olmarker K, Olmarker K. Indomethacin blocks the nucleus pulposus-induced effects on nerve root function. An experimental study in dogs with assessment of nerve conduction and blood flow following experimental disc herniation. (0940-6719 (Print)).

21. Bourinbaiar AS, Lee-Huang S. The non-steroidal anti-inflammatory drug, indomethacin, as an inhibitor of HIV replication. (0014-5793 (Print)).

22. De Clercq E, Li G. Approved Antiviral Drugs over the Past 50 Years. (1098-6618 (Electronic)).

23. Vane JR, Botting RM. Mechanism of action of anti-inflammatory drugs. Adv Exp Med Biol. 1997;433:131–138.

24. García MA, Gil J Fau - Ventoso I, Ventoso I Fau - Guerra S, et al. Impact of protein kinase PKR in cell biology: from antiviral to antiproliferative action. (1092-2172 (Print)).

25. Dabo S, Meurs EF. dsRNA-dependent protein kinase PKR and its role in stress, signaling and HCV infection. (1999-4915 (Electronic)).

26. www.plexresearch.com

27. Liedtke AJ, Adeniji AO, Chen M, et al. Development of potent and selective indomethacin analogues for the inhibition of AKR1C3 (Type 5 17β-hydroxysteroid dehydrogenase/prostaglandin F synthase) in castrate-resistant prostate cancer. Journal of medicinal chemistry. 2013;56(6):2429–2446.

28. Jamieson SM, Brooke DG, Heinrich D, et al. 3-(3,4-Dihydroisoquinolin-2(1H)-ylsulfonyl)benzoic Acids: highly potent and selective inhibitors of the type 5 17-β-hydroxysteroid dehydrogenase AKR1C3. Journal of medicinal chemistry. 2012;55(17):7746–7758.

29. Endo S, Matsunaga T, Kanamori A, et al. Selective inhibition of human type-5 17β-hydroxysteroid dehydrogenase (AKR1C3) by baccharin, a component of Brazilian propolis. Journal of natural products. 2012;75(4):716–721.

30. Cerelli MJ, Curtis DL, Dunn JP, Nelson PH, Peak TM, Waterbury LD. Antiinflammatory and aldose reductase inhibitory activity of some tricyclic arylacetic acids. Journal of medicinal chemistry. 1986;29(11):2347–2351.

31. Acton JJ, 3rd, Black RM, Jones AB, et al. Benzoyl 2-methyl indoles as selective PPARgamma modulators. Bioorganic & medicinal chemistry letters. 2005;15(2):357–362.

32. Romeiro NC, Sant’Anna CM, Lima LM, Fraga CA, Barreiro EJ. NSAIDs revisited: putative molecular basis of their interactions with peroxisome proliferator-activated gamma receptor (PPARgamma). European journal of medicinal chemistry. 2008;43(9):1918–1925.

33. Lee JY, Kim JK, Cho MC, et al. Cytotoxic flavonoids as agonists of peroxisome proliferator-activated receptor gamma on human cervical and prostate cancer cells. Journal of natural products. 2010;73(7):1261–1265.

34. Gallant M, Dufresne C, Gareau Y, et al. New class of potent ligands for the human peripheral cannabinoid receptor. Bioorganic & medicinal chemistry letters. 1996;6(19):2263–2268.

35. Nguyen TA, Smith BRC, Tate MD, et al. SIDT2 Transports Extracellular dsRNA into the Cytoplasm for Innate Immune Recognition. Immunity. 2017;47(3):498–509.e496.

36. Ballekova J, Soltesova-Prnova M Fau - Majekova M, Majekova M Fau - Stefek M, Stefek M. Does inhibition of aldose reductase contribute to the anti-inflammatory action of setipiprant? (1802-9973 (Electronic)).

37. Ramana KV, Srivastava SK. Aldose reductase: a novel therapeutic target for inflammatory pathologies. (1878-5875 (Electronic)).

38. Jiang C, Ting AT, Seed B. PPAR-gamma agonists inhibit production of monocyte inflammatory cytokines. Nature. 1998;391(6662):82–86.

39. Heming M, Gran S, Jauch SL, et al. Peroxisome Proliferator-Activated Receptor-γ Modulates the Response of Macrophages to Lipopolysaccharide and Glucocorticoids. Frontiers in immunology. 2018;9:893.

40. Kaplan BLF, Rockwell CE, Kaminski NE. Evidence for cannabinoid receptor-dependent and -independent mechanisms of action in leukocytes. J Pharmacol Exp Ther. 2003;306(3):1077–1085.

41. Rom S, Persidsky Y. Cannabinoid receptor 2: potential role in immunomodulation and neuroinflammation. J Neuroimmune Pharmacol. 2013;8(3):608–620.

42. Miller AM, Stella N. CB2 receptor-mediated migration of immune cells: it can go either way. Br J Pharmacol. 2008;153(2):299–308.

43. Kurihara R, Tohyama Y, Matsusaka S, et al. Effects of peripheral cannabinoid receptor ligands on motility and polarization in neutrophil-like HL60 cells and human neutrophils. The Journal of biological chemistry. 2006;281(18):12908–12918.

44. Klein TW, Cabral GA. Cannabinoid-induced immune suppression and modulation of antigen-presenting cells. J Neuroimmune Pharmacol. 2006;1(1):50–64.

45. Tanaka T, Narazaki M, Kishimoto T. IL-6 in inflammation, immunity, and disease. (1943-0264 (Electronic)).

46. Zampronio AR, Melo Mc Fau - Silva CA, Silva Ca Fau - Pelá IR, Pelá Ir Fau - Hopkins SJ, Hopkins Sj Fau - Souza GE, Souza GE. A pre-formed Pyrogenic Factor Released by Lipopolysaccharide Stimulated Macrophages. (0962-9351 (Print)).

47. Chen L, Liu HG, Liu W, et al. Analysis of clinical features of 29 patients with 2019 novel coronavirus pneumonia. Zhonghua Jie He He Hu Xi Za Zhi. 2020;43(0):E005–E005.

48. Loppnow H, Zhang L, Buerke M, et al. Statins potently reduce the cytokine-mediated IL-6 release in SMC/MNC cocultures. J Cell Mol Med. 2011;15(4):994–1004.

49. Gordon DE, Jang GM, Bouhaddou M, et al. 2020.

50. Nguyen TA, Smith BRC, Tate MD, et al. SIDT2 Transports Extracellular dsRNA into the Cytoplasm for Innate Immune Recognition. (1097-4180 (Electronic)).

51. Vane JR. Inhibition of prostaglandin synthesis as a mechanism of action for aspirin-like drugs. Nature: New biology. 1971;231(25):232–235.

52. Chan JF, Kok KH, Zhu Z, et al. Genomic characterization of the 2019 novel human-pathogenic coronavirus isolated from a patient with atypical pneumonia after visiting Wuhan. Emerging microbes & infections. 2020;9(1):221–236.

53. Maier HJ, Bickerton E, Britton P. Preface. Coronaviruses. Methods in molecular biology. 2015;1282:v.

54. Taytard A, Daures JP, Arsac P, et al. [Management of lower respiratory tract infections by general practitioners in France]. Revue des maladies respiratoires. 2001;18(2):163–170.

55. Raherison C, Peray P, Poirier R, et al. Management of lower respiratory tract infections by French general practitioners: the AIR II study. Analyse Infections Respiratoires. The European respiratory journal. 2002;19(2):314–319.

56. Raherison C, Poirier R, Daurès JP, et al. Lower respiratory tract infections in adults: non-antibiotic prescriptions by GPs. Respiratory medicine. 2003;97(9):995–1000.

57. Vane JR, Botting RM. Mechanism of action of anti-inflammatory drugs. (0065-2598 (Print)).

